# A biosecurity baseline for transboundary management of marine biological invasions in the ROPME Sea Area

**DOI:** 10.64898/2026.03.13.711635

**Authors:** Lorenzo Vilizzi, Ali Mansoor Abbas, Maryam A. Mubarak, Mohammad Hadi Alavi, Mahtab Shojaei, Daryoush Moghaddas, Hossein Rahmani, Ahmed A. R. Albu Salih, Mohammed F. A. Al-Khayyat, Abbas J. Al-Faisal, Aisha F. Al-Marhoun, Ali H. Abdulhussain, Jumana Alkhamees, Qusaie E. Karam, Wafa’a Behbehani, Mohammed Al Rezaiqi, Muna Al Tarshi, Salman F. Al-Dosari, Amal M. Al Jamaei, Mohammed Elnaiem A. El Mahdi, Abdirahman A. Mohamed, Eman I. Sabbagh, Ebrahim A. Jamali, Nahla Mehzoud, Obaid A. H. Al Shamsi, Zainab Al-Wazzan

**Author notes:** Corresponding author. University of Lodz, Faculty of Biology and Environmental Protection, Department of Ecology and Vertebrate Zoology, Lodz, Poland (L. Vilizzi). E-mail addresses:* (A. M. Abbas), (M. A. Mubarak), (M. H. Alavi), (M. Shojaei), (D. Moghaddas), (H. Rahmani), (A. A. R. Albu Salih), (M. F. A. Al-Khayyat), (A. J. Al-Faisal), (A. F. Al-Marhoun), (A. H. Abdulhussain), (J. Alkhamees), (Q. E. Karam), (W. Behbehani), (M. Al Rezaiqi), (M. Al Tarshi), (S. F. Al-Dosari), (A. M. Al Jamaei), (M. E. A. El Mahdi), (A. A. Mohamed), (E. I. Sabbagh), (E. A. Jamali), (N. Mezhoud), (O. A. H. Al Shamsi), (Z. Al-Wazzan).

## Abstract

Marine and brackish-water ecosystems are increasingly degraded by cumulative human pressures, with biological invasions representing a major driver of biodiversity loss, ecosystem disruption, and socio-economic impacts. Effective management requires regionally harmonized and scientifically robust baselines capable of supporting coordinated transboundary decision-making. Here we present the first consolidated marine biosecurity baseline for the Regional Organization for the Protection of the Marine Environment (ROPME) Sea Area, a transboundary region characterized by extreme environmental conditions and increasing biosecurity pressure. A total of 192 species (123 extant and 69 horizon), including birds, fishes, tunicates, invertebrates, plants, and chromists, were systematically reviewed, taxonomically validated, and cross-checked against major databases and Member State inputs. Re-evaluation of a previous regional screening revealed substantial inconsistencies, with 24 species (≈18%) requiring status correction or exclusion. The resulting consolidated inventory comprised 130 validated retained species supplemented by 62 additional taxa. Extant species were classified according to biogeographic origin and impact status, whereas horizon species were evaluated based on introduction pathways, environmental suitability, and projected climate trends. Risk screening under current and projected climate conditions identified 39 extant species as very high risk, providing an operational basis for progression to full risk assessment and coordinated regional biosecurity management.

## 1 Introduction

Globally, marine and brackish-water ecosystems are undergoing accelerating degradation under cumulative human-induced pressures, underscoring the need for strengthened conservation and adaptive management, particularly in environmentally fragile regions (Halpern et al., 2019; Duarte et al., 2020; Sala et al., 2021; Campagne et al., 2023; Vargas-Fonseca et al., 2024). Among these pressures, biological invasions are widely recognized as a primary driver of biodiversity loss, ecosystem restructuring, and socio-economic impact (Early et al., 2016; Seebens et al., 2017, 2021; Diagne et al., 2021; Roy et al., 2024). In line with international policy frameworks, including the Convention on Biological Diversity (https://www.cbd.int/), effective management of marine invasive species prioritizes prevention, followed by early detection, rapid response, pathway management, and containment where eradication is infeasible (Roy et al., 2019; Hulme, 2020; O’Shaughnessy et al., 2023). This is because, once established, marine non-native species are notoriously difficult to eradicate (Williams and Grosholz, 2008; Werschkun et al., 2014). Achieving these objectives requires regionally harmonized baselines that are scientifically robust, regularly updated, and institutionally coordinated across jurisdictions (e.g. Ojaveer et al., 2018; Tiralongo et al., 2022).

These challenges are particularly acute in the Regional Organization for the Protection of the Marine Environment (ROPME) Sea Area (RSA), which comprises the Gulf, the Sea of Oman, and adjacent parts of the Arabian Sea. The RSA includes the Inner, Middle, and Outer subregions and is bordered by Bahrain, Iran, Iraq, Kuwait, Oman, Qatar, Saudi Arabia, and the United Arab Emirates (UAE), which constitute the ROPME Member States (Van Lavieren and Klaus, 2013; Bailey and Munawar, 2015) (Fig. 1).

**Figure 1.**
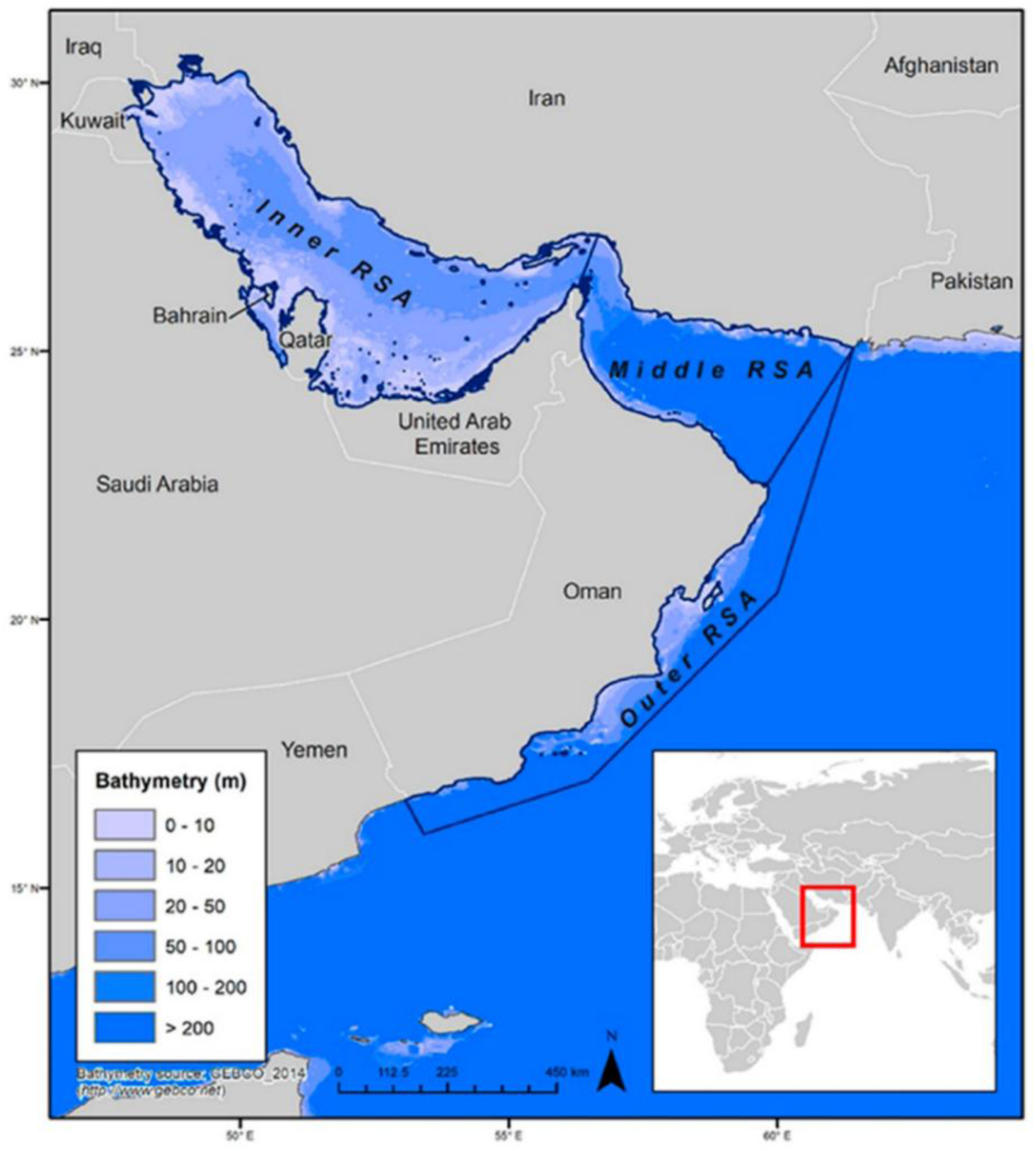
Map of the study and risk assessment area: the Regional Organization for the Protection of the Marine Environment (ROPME) Sea Area (RSA) encompassing the Gulf, the Sea of Oman, and parts of the Arabian Sea, and subdivided into Inner, Middle, and Outer regions.

The RSA represents one of the most environmentally extreme marine systems globally, characterized by naturally low species richness and communities living close to physiological tolerance limits, conditions that confer intrinsic sensitivity to disturbance (Lincoln et al., 2021). Rapid coastal urbanization, port expansion, desalination, aquaculture growth, and population increase have intensified cumulative pressures in recent decades (Sale et al., 2011; Riegl et al., 2012; Burt, 2014; Bailey and Munawar, 2015), while recurrent harmful algal blooms (HABs) linked to eutrophication, climatic variability, and maritime activity have caused ecological damage and operational disruption of desalination infrastructure (Al-Yamani et al., 2012). Oceanographic constraints, including shallow depth, restricted circulation, and limited exchange with the Indian Ocean through the Strait of Hormuz, further amplify system-wide vulnerability (Ezam et al., 2010; Al-Said et al., 2018). Although mangroves, seagrasses, and thermally tolerant coral assemblages persist, ecological recovery capacity remains limited (Rezai et al., 2004). Concurrently, the RSA functions as one of the world’s busiest maritime corridors (Bayani, 2016; Khazali, 2021), where intensive shipping, coastal infrastructure development, and aquaculture elevate propagule pressure through ballast water, hull fouling, vessel traffic, stock transfers, and aquarium releases (Sale et al., 2011). Rising sea temperatures, altered salinity regimes, desalination brine discharge, and nutrient enrichment may further modify habitat suitability and competitive dynamics, compounding the interaction between climate stress and biological invasion risk across the region (Maltby et al., 2022; Paparella et al., 2022).

Despite increasing awareness of invasion risk in the RSA, a fully validated and regionally endorsed biosecurity reference framework has remained lacking. Recognizing these pressures, Clarke et al. (2020) conducted the first structured screening of extant and horizon marine and brackish-water species for the Gulf and Sea of Oman, covering the Inner and Middle RSA. That study represented an important regional milestone, providing an initial risk-based inventory to inform biosecurity awareness. However, it did not incorporate systematic cross-validation of species status across all ROPME Member States, full taxonomic verification, updated distributional data, or harmonized categorization consistent with contemporary invasion-science frameworks (Blackburn et al., 2011; Essl et al., 2019). These limitations have constrained its direct operational use for coordinated transboundary biosecurity planning.

The aim of this study was to provide a regional biosecurity baseline to establish the first regionally harmonized and scientifically consolidated marine inventory for the RSA. Specifically, we (i) reviewed, corrected, and updated the list of extant and horizon species previously screened, incorporating verified distributional records across all ROPME Member States and newly available evidence; (ii) categorized extant species according to contemporary invasion-science frameworks; (iii) conducted risk identification of extant and horizon species under current and projected climate conditions; (iv) calibrated and validated the resulting dataset and ranked species by organismal group according to their level of risk to ensure transparent evidence quality and prioritization for subsequent full risk assessment and coordinated management. Collectively, this process establishes a regionally harmonized operational biosecurity baseline for the RSA.

## 2 Methods

### 2.1 Species selection

The dual-status approach of including both extant and horizon species in risk identification studies (Vilizzi et al., 2022a) was adopted for species selection in the RSA (risk assessment area). Extant species are those currently recorded in the RSA either in the wild or under confinement in human-controlled facilities, whereas horizon species are those not yet recorded but considered likely to arrive, establish, or generate ecological or socio-economic impacts based on known introduction pathways, environmental suitability, regional climate trends, and documented invasion histories elsewhere. The taxonomic scope of the assessment was intentionally comprehensive and included all marine and brackish-water organism groups relevant to invasion risk. Although the study primarily concerns zoological taxa, aquatic macrophytes and other marine plants were also retained because habitat-forming species can play a significant role in invasion dynamics and ecosystem alteration in coastal environments, particularly in semi-enclosed seas. Plant taxa represent only 19 of the 192 screened species and were evaluated using the same risk-screening methodology applied to the zoological taxa, ensuring that the resulting inventory reflects the full spectrum of organisms relevant to regional marine biosecurity.

To ensure accurate and scientifically robust risk identification, we re-evaluated the assignments into extant and horizon of the 136 species originally screened by Clarke et al. (2020). We retained 130 species from the original list (70 extant and 60 horizon) and subjected them to updated screening (see Section 2.3). We identified and screened an additional 62 species, of which 53 extant and nine horizon, through a systematic search of regional non-native species databases, technical reports, and peer-reviewed literature (Supplementary Material 1: available at https://doi.org/10.6084/m9.figshare.31654384). This approach ensured a comprehensive, current, and regionally validated inventory of extant species in the RSA while maintaining a proportionate and strategically targeted expansion of the horizon species list originally developed by Clarke et al. (2020). The final validated list included 192 species in total of which 123 extant and 69 horizon (Table S1).

### 2.2 Taxonomic validation and geographic verification

We validated the taxonomic nomenclature of all 192 species against the World Register of Marine Species (WoRMS: https://www.marinespecies.org/). For extant species, we verified geographic occurrence at the level of individual Member States using national non-native species databases, Global Register of Introduced and Invasive Species (GRIIS: Pagad et al., 2018) country lists, official technical reports, and peer-reviewed literature. For Iran and Saudi Arabia, we excluded records referring exclusively to the Caspian Sea and Red Sea, respectively, as these fall outside the defined marine and brackish-water boundaries of the RSA. For horizon species, we verified absence of confirmed records within the RSA while documenting invasion history, environmental suitability, and introduction pathways using peer-reviewed literature and global invasive species databases.

### 2.3 Extant species

#### 2.3.1 Listing

For the 123 validated extant species, we compiled information on their listing and status across the ROPME Member States using regional and global non-native species databases, official national reports, and peer-reviewed literature. We sourced occurrence and status information from global and regional non-native species databases (Seebens et al., 2020; Briski et al., 2024), country-level GRIIS lists (Iran: Sohrabi and Pagad, 2024; Iraq: Haloob et al., 2020; Kuwait: Pagad, 2022; Oman: Patzelt and Pagad, 2020; Qatar: Elazazi et al., 2020; Saudi Arabia: Pandalayil et al., 2020; United Arab Emirates: Al Ali et al., 2020), official Member State reports (Iran: Department of Environment Islamic Republic of Iran, 2025; Iraq: Ministry of Health and Environment, Republic of Iraq, 2018; Kuwait: Al-Yamani, 2021; United Arab Emirates: UAE MOCCAE, 2022), and global invasive species information systems including Avibase (Lepage et al., 2014), the Global Biodiversity Information Facility (GBIF: https://www.gbif.org/), the Global Invasive Species Database (GISD: https://www.iucngisd.org/gisd/), the Smithsonian Environmental Research Center’s National Estuarine and Marine Exotic Species Information System (NEMESIS: https://invasions.si.edu/nemesis/), and the Ocean Biodiversity Information System (OBIS: https://obis.org/).

#### 2.3.2 Categorization

To ensure consistent classification and ecological interpretation, we categorized each of the 123 extant species according to biogeographic origin, dispersal history, and documented ecological impact. For each species, we conducted a comprehensive review of peer-reviewed literature and official technical reports, with particular emphasis on the Gulf and its Member States. We integrated this information with data compiled from non-native species databases and national reports (see Section 2.3.1).

We performed status assignment at the level of individual Member States. Accordingly, even when a species occurred in multiple Member States, we determined its status only in those jurisdictions where its presence and ecological attributes indicative of invasiveness or harmful impact were explicitly documented in published or verified sources. For example, dinoflagellate species reported broadly across the Gulf were categorized only in those Member States where they were identified as ecologically or socio-economically impactful. Conversely, species listed solely in faunal or floral inventories without evidence of ecological impact or invasion status were not categorized as invasive or nuisance.

This categorization framework draws on established invasion-science theory (Blackburn et al., 2011; Essl et al., 2019) and recent advances in terminology refinement to support management and policy applications (Soto et al. 2024; Vilizzi et al., 2025b, 2026). We distinguished four primary origin categories: (i) native species, historically part of the regional biota within a given Member State; (ii) neonative species, whose natural dispersal has been facilitated by anthropogenic environmental change; (iii) cryptogenic species, of uncertain biogeographic origin; and (iv) non-native species, introduced through direct or indirect human activity. For native, neonative, and cryptogenic species, we additionally identified nuisance forms where species currently exert locally harmful ecological or socio-economic impacts under contemporary environmental conditions. We also distinguished non-native captive species, defined as organisms present exclusively in confinement (e.g. aquaculture facilities) without confirmed establishment in the wild. We structured the categorization to distinguish explicitly between biogeographic origin and documented ecological impact (Supplementary Material 1: available at https://doi.org/10.6084/m9.figshare.31654384). Under this framework, a species may hold different status designations among Member States (e.g. nuisance native in one jurisdiction and neonative in another), reflecting documented local ecological conditions.

### 2.4 Risk identification

We employed the Aquatic Species Invasiveness Screening Kit (AS-ISK) v2.4.1 (Copp et al., 2016c; Vilizzi et al., 2025a; available at https://tinyurl.com/ISK-toolkits), a taxon-generic decision-support tool comprising 55 questions of which 49 constitute the Basic Risk Assessment (BRA) and six the Climate Change Assessment (CCA). The BRA evaluates species biogeography, invasion history, biology, and ecology, while the CCA assesses how projected climatic conditions may modify risks of introduction, establishment, dispersal, and impact.

Twenty-six assessors, all co-authors of this study, conducted the screenings. Twenty-five assessors updated the screenings for the 130 species retained from Clarke et al. (2020), with each assessor evaluating between one and 12 species; 104 species were updated by a single assessor, 21 by two assessors jointly, and five by four assessors jointly. Twenty-two assessors screened the 62 additional species, with 52 species evaluated by a single assessor and 10 jointly by two assessors. All assessors possess expertise in marine and brackish-water ecology in the RSA and received structured training in AS-ISK application during a ROPME regional workshop held in Kuwait in January 2025, led by the senior author. The senior author supervised the full screening process, reviewed all 192 assessments for methodological consistency, and coordinated interpretation of scoring criteria following established AS-ISK guidance (Vilizzi and Piria, 2022), thereby ensuring a consensus-based application across taxa and Member States (Vilizzi et al., 2022a).

We followed the standard AS-ISK protocol (Vilizzi and Piria, 2022; Vilizzi et al., 2022a, 2022b). For each question, assessors provided a response, a confidence level, and supporting justification. The tool generates two numerical outputs: the BRA score and the combined BRA+CCA score. Scores < 1 indicate low risk, whereas scores ≥ 1 indicate medium or high risk. Differentiation between medium- and high-risk categories requires calibration through Receiver Operating Characteristic (ROC) curve analysis based on *a priori* classification of species as invasive or non-invasive (Vilizzi et al., 2022a; Table S2). Calibration requires at least 15–20 species with balanced representation of *a priori* invasive and non-invasive. This condition was satisfied for fishes, invertebrates, plants, and chromists, but not for birds and tunicates. For the latter groups, we applied generalized thresholds of 25.5 and 22.5, respectively, following Vilizzi et al. (2021). Because brackish subsets were insufficiently large for separate calibration, we pooled marine and brackish species within fishes, invertebrates, and plants for threshold estimation. For species classified as high risk, we defined an additional ‘very high risk’ category (score ≥ 50) to support operational prioritization for comprehensive risk assessment. Such assessments evaluate risks of introduction, establishment of self-sustaining populations, secondary spread within the assessment area, and ecological, socio-economic, or disease-related impacts.

Implementation of ROC curve analysis followed the standard protocol (Vilizzi et al., 2022a). True/false positive/negative outcome classification was not applied to medium-risk species because their progression to comprehensive risk assessment depends on policy priorities and resource availability. AUC values were interpreted according to Hosmer et al. (2013). We performed ROC analyses using the pROC package (Robin et al., 2011) in R v4.5.1 (R Core Team, 2025) with optimal thresholds determined by maximizing Youden’s *J* statistic while retaining the default threshold of 1 to distinguish low- from medium-risk species. We computed confidence levels for each response and resulting confidence factor (CF) as per Vilizzi et al. (2022a): CF_Total_, CF_BRA_, and CF_CCA_ using all 55 questions, the 49 BRA questions, and the six CCA questions, respectively. We tested differences in CF between BRA and BRA+CCA components using permutational ANOVA on normalized data, based on Bray–Curtis dissimilarity and 9,999 unrestricted permutations, with significance evaluated at α = 0.05.

## 3 Results

### 3.1 Species selection

Of the 136 species originally screened, 56 were classified as extant and 80 as horizon for the RSA. Following re-evaluation of the status of each of these species, several notable discrepancies were observed (Table S1).

Of the 56 species classified as extant, 54 were confirmed as such whereas two were found to be horizon: scaly tunicate *Microcosmus squamiger* and *Prorocentrum mexicanum*. Of the 80 species classified as horizon, 58 were confirmed as such whereas of the other 22 species:

- 16 were found to be extant: *Acartia (Acanthacartia) tonsa*, pile worm *Alitta succinea*, Asian date mussel *Arcuatula senhousia*, Pharaoh’s mussel *Brachidontes pharaonis*, green sea fingers *Codium fragile fragile*, European seabass *Dicentrarchus labrax*, Chinese mitten crab *Eriocheir sinensis*, ragworm *Hediste diversicolor*, Chinese freshwater mussel *Limnoperna fortunei*, whiteleg shrimp *Penaeus (Litopenaeus) vannamei*, kuruma prawn *Penaeus (Marsupenaeus) japonicus*, brown mussel *Perna perna*, New Zealand mudsnail *Potamopyrgus antipodarum*, golden alga *Prymnesium parvum*, gulf weed *Sargassum fluitans*, red-rust bryozoan *Watersipora subtorquata*;
- five were found to be native (and excluded from analysis): yellowfin seabream *Acanthopagrus latus*, snowflake coral *Carijoa riisei*, killer algae *Caulerpa taxifolia*, red lionfish *Pterois volitans*, nomad jellyfish *Rhopilema nomadica*;
- one was found to be only freshwater (and excluded from analysis): Texas cichlid *Herichthys cyanoguttatus*, hence of no relevance to the habitats of the RSA including its coastal and estuarine areas.

For the horizon to extant species, literature evidence indicated that only *Arcuatula senhousia* and *Limnoperna fortunei* entered the risk assessment area after 2019 (i.e. the year prior to publication of Clarke et al., 2020), whereas all other species were already present (hence, extant) by then, including *Penaeus (Litopenaeus) vannamei*, although as a non-native captive species. As per *Acanthopagrus latus*, originally classified as horizon, this marine fish is part of a species complex that occurs naturally in the Gulf as *Acanthopagrus arabicus* sp. nov. and *Acanthopagrus sheim* (Iwatsuki, 2013).

Overall, revision of the original species list revealed that the status in the RSA of 24 species (≈ 18%) had been incorrectly determined. As a result of the exclusion of six species, 130 species in total, comprising 70 extant and 60 horizon, as per the status re-evaluation, were retained from that list. The original list was then augmented with 62 additional species of which 53 extant and nine horizon (Table S1). This resulted in 192 species in total, of which 123 extant and 69 horizon, that were included in the present study for risk identification. The species comprised two birds (both extant), 37 fishes (13 extant and 24 horizon; 17 brackish and 20 marine), 14 tunicates (13 extant and six horizon), 74 invertebrates (43 extant and 31 horizon; five brackish and 69 marine), 19 plants (13 extant and six horizon; three brackish, 16 marine), and 46 chromists (44 extant and two horizon; all marine) (Table S1; Supplementary Material 2: available at https://doi.org/10.6084/m9.figshare.31654384).

### 3.2 Extant species

Of the 123 extant species identified for the RSA, 102 were recorded in at least one non-native species database or official report. The number of listed species varied markedly among Member States: nine in Bahrain, 26 in Iran, 19 in Iraq, 35 in Kuwait, 48 in Oman, six in Qatar, 16 in Saudi Arabia, and 27 in the United Arab Emirates (Table S3).

Comparable variation across Member States was observed when species were classified across the eight status categories (Table 1; Table S4), as well as into broader origin–impact classes (Fig. 2). Whereas the majority of Member States exhibited a higher proportional dominance of non-native species, Kuwait and especially Oman were characterized by a substantial nuisance component. Differences were also evident at the subregional scale, with 114 species recorded in the Inner RSA, 58 in the Middle RSA, and 12 in the Outer RSA (Table S4).

**Figure 2.**
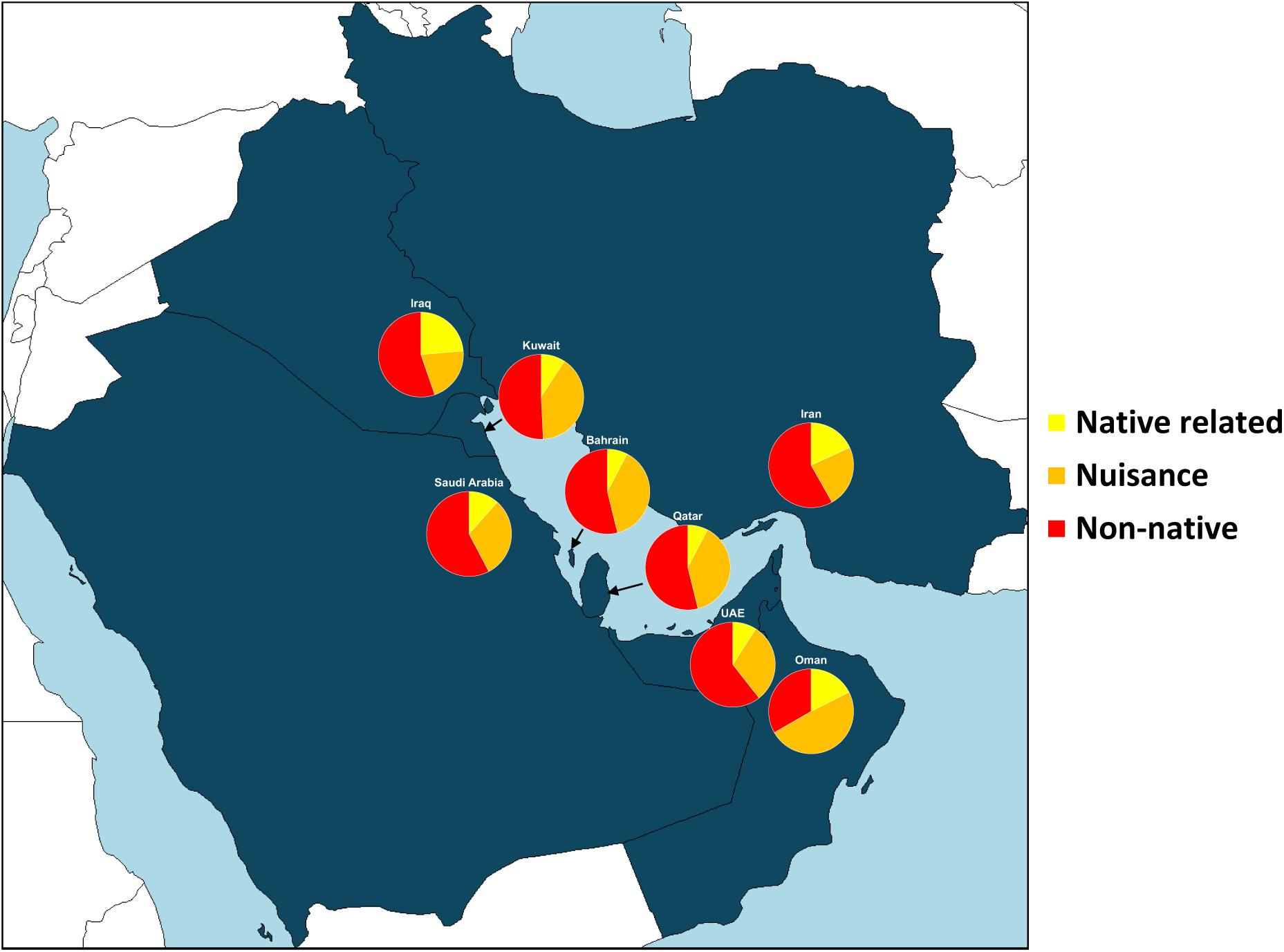
Relative composition of extant species across ROPME Member States grouped into broad origin–impact classes: native-related (native, neonative, cryptogenic), nuisance (nuisance native, nuisance neonative, nuisance cryptogenic), and non-native (non-native captive and non-native) (see Table 1). Values represent proportions within each Member State based on the validated extant species inventory (*n* = 123).

**Table 1.** Number (*n*) and percentage (%) of extant species (*n* = 123) recorded in each Member State of the Regional Organisation for the Protection of the Marine Environment (ROPME) Sea Area classified according to biogeographic origin and impact status. Percentages are calculated relative to the total number of extant species recorded within each Member State. Status assignment was conducted at the Member State level; consequently, a given species may hold different status designations across jurisdictions depending on documented origin and impact. See also Table S4.

| Status | Bahrain |  | Iran |  | Iraq |  | Kuwait |  | Oman |  | Qatar |  | Saudi Arabia |  | UAE |  |
| --- | --- | --- | --- | --- | --- | --- | --- | --- | --- | --- | --- | --- | --- | --- | --- | --- |
|  | *n* | % | *n* | % | *n* | % | *n* | % | *n* | % | *n* | % | *n* | % | *n* | % |
| Native | 0 | 0.0 | 2 | 3.7 | 0 | 0.0 | 0 | 0.0 | 6 | 10.5 | 0 | 0.0 | 0 | 0.0 | 0 | 0.0 |
| Neonative | 1 | 7.7 | 4 | 7.4 | 7 | 18.9 | 4 | 6.2 | 2 | 3.5 | 1 | 7.7 | 3 | 11.5 | 2 | 5.9 |
| Cryptogenic | 1 | 7.7 | 3 | 5.6 | 1 | 2.7 | 2 | 3.1 | 2 | 3.5 | 0 | 0.0 | 0 | 0.0 | 2 | 5.9 |
| Nuisance native | 0 | 0.0 | 9 | 16.7 | 5 | 13.5 | 20 | 30.8 | 23 | 40.4 | 0 | 0.0 | 1 | 3.8 | 6 | 17.6 |
| Nuisance neonative | 2 | 15.4 | 3 | 5.6 | 3 | 8.1 | 5 | 7.7 | 4 | 7.0 | 3 | 23.1 | 5 | 19.2 | 3 | 8.8 |
| Nuisance cryptogenic | 3 | 23.1 | 1 | 1.9 | 0 | 0.0 | 1 | 1.5 | 1 | 1.8 | 2 | 15.4 | 2 | 7.7 | 1 | 2.9 |
| Non-native captive | 1 | 7.7 | 4 | 7.4 | 1 | 2.7 | 5 | 7.7 | 4 | 7.0 | 0 | 0.0 | 5 | 19.2 | 7 | 20.6 |
| Non-native | 5 | 38.5 | 28 | 51.9 | 20 | 54.1 | 28 | 43.1 | 15 | 26.3 | 7 | 53.8 | 10 | 38.5 | 13 | 38.2 |
| **Total** | **13** |  | **54** |  | **37** |  | **65** |  | **57** |  | **13** |  | **26** |  | **34** |  |

### 3.3 Risk identification

Application of AS-ISK v2.4.1 across the 192 species generated calibrated and confidence-weighted invasion-risk profiles for the RSA under both baseline (BRA) and climate-adjusted (BRA+CCA) scenarios. Complete AS-ISK v2.4.1 reports, including component scores and confidence values under BRA and BRA+CCA scenarios, are provided in Supplementary Material 3 (available at https://doi.org/10.6084/m9.figshare.31654384).

Across all species, mean confidence values were 0.690 ± 0.008 SE (CF_Total_), 0.701 ± 0.007 SE (CF_BRA_), and 0.601 ± 0.013 SE (CF_CCA_) (Table S5). Mean CF_BRA_ was significantly higher than mean CF_CCA_ (permutational ANOVA, *F*^#^_1,382_ = 41.84, *P*^#^ < 0.001), where # denotes permutational values.

#### 3.3.1 Organismal groups

*Birds* – The generalized threshold of 25.5 was applied for calibration (Fig. 3; Table S5). Based on both BRA and BRA+CCA scores, one species was classified as high risk (extant African sacred ibis *Threskiornis aethiopicus*) and one as medium risk (extant greater flamingo *Phoenicopterus ruber*), with both species extant in the RSA. Using the ≥ 50 cut-off, *T. aethiopicus* ranked as very high risk under BRA+CCA. Climate-change adjustment increased the BRA score for *T. aethiopicus* and decreased it for *P. ruber*.

**Figure 3.**
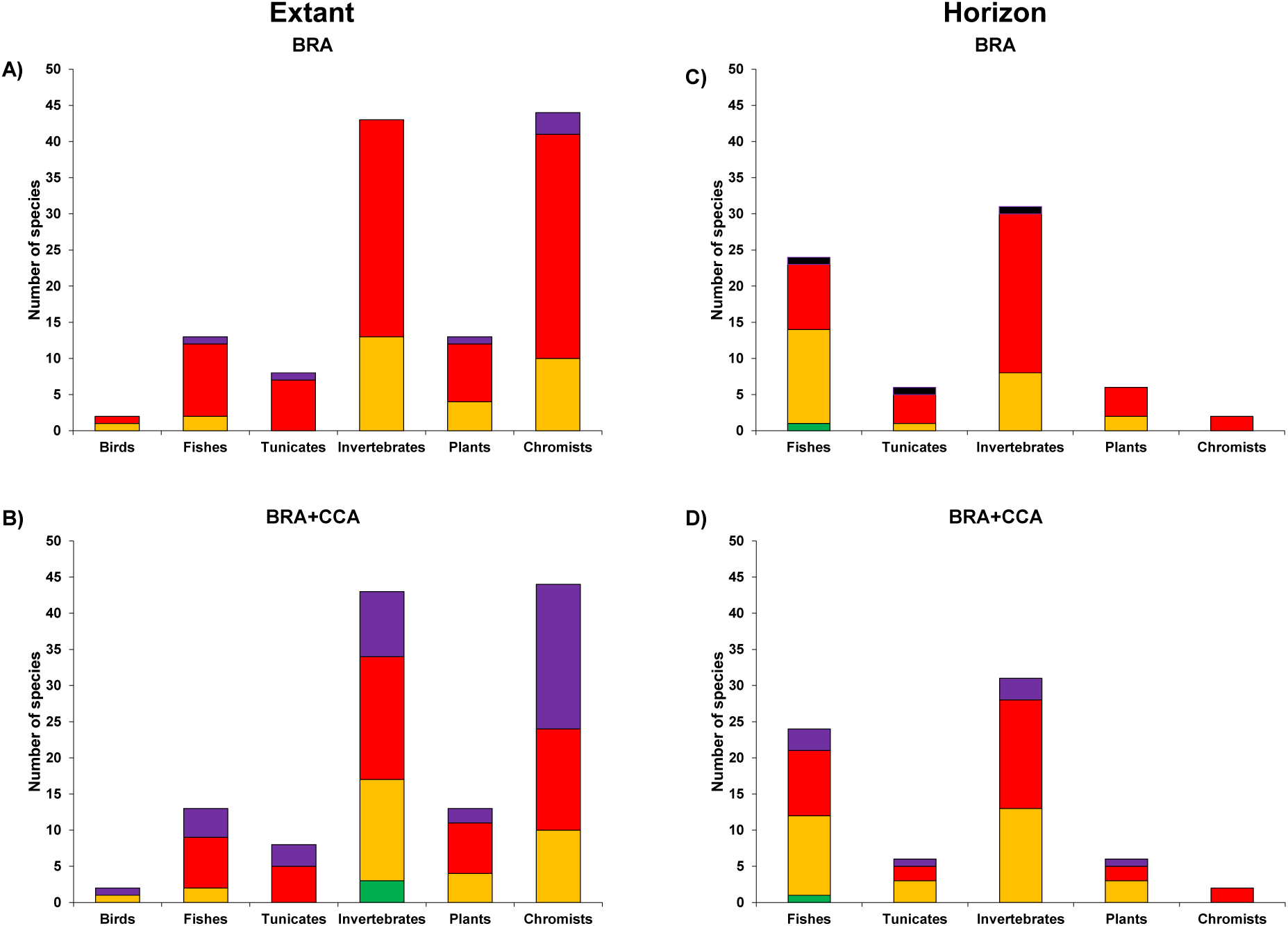
Number of extant and horizon species by organismal group assessed with the Aquatic Species Invasiveness Screening Kit for the RSA. For each group, the proportion of species posing a low (green), medium (orange), high (red), and very high risk level is provided for (a, c) the Basic Risk Assessment (BRA), and (b, d) the BRA + Climate Change Assessment (CCA). Group-specific thresholds to distinguish between medium and high risk species are given in Table S5, with distinction between low and medium risk species based on a fixed threshold of 1, and between high-risk and very high-risk spocies on an *ad hoc* threshold of 50.

*Fishes* – ROC analysis yielded an AUC of 0.6838 (95% CI: 0.5082–0.8594) and a calibrated threshold of 25.5 (Fig. 3; Table S5). Based on BRA scores, 21 of 37 species (56.8%) were classified as high risk, 15 (40.5%) as medium risk, and one (2.7%) as low risk (horizon big-scale pomfret *Taractichthys longipinnis*). Under BRA+CCA, 23 species (62.2%) were classified as high risk, 13 (35.1%) as medium risk, and one (2.7%) remained low risk (*T. longipinnis*). Using the ≥ 50 cut-off, there were one very high-risk horizon species under BRA (Mayan cichlid *Mayaheros urophthalmus*), one extant under both BRA and BRA+CCA (Mozambique tilapia *Oreochromis mossambicus*), and five additional species under BRA+CCA (extant redbelly tilapia *Coptodon zillii* and sailfin molly *Poecilia latipinna*; horizon spinycheek grouper *Epinephelus fuscoguttatus*, Wami tilapia *Oreochromis urolepis*, blackchin tilapia *Sarotherodon melanotheron*). Climate-change adjustment increased scores for 18 species (48.7%), resulted in no change for three (8.1%), and decreased scores for 16 (43.2%).

*Tunicates* – The generalized threshold of 22.5 was applied (Fig. 3; Table S5). Based on BRA scores, 13 of 14 species (92.9%) were classified as high risk, and one (7.1%) as medium risk and *a priori* invasive. Under BRA+CCA, 11 species (78.6%) were classified as high risk, and three (21.4%) as medium risk. Using the ≥ 50 cut-off, there were two very high-risk species under both BRA and BRA+CCA (extant *Didemnum psammatodes*, horizon *Polyandrocarpa zorritensis*) and two additional extant species under BRA+CCA (black colonial tunicate *Botrylloides niger*, *Symplegma brakenhielmi*). Climate-change adjustment increased scores for six species (42.9%), resulted in no change for two (14.2%), and decreased scores for six (42.9%).

*Invertebrates* – ROC analysis yielded an AUC of 0.7991 (95% CI: 0.6558–0.9424) and a calibrated threshold of 23.25 (Fig. 3; Table S5). Based on BRA scores, 53 of 74 species (71.6%) were classified as high risk, and 21 (28.4%) as medium risk. Under BRA+CCA, 44 species (59.5%) were classified as high risk, 27 (36.5%) as medium risk, and three (4.1%) extant species as low risk and true negatives (dark doto *Doto kya*, opossum shrimp *Rhopalophthalmus tattersallae*, white-crust cuthona *Tenellia albocrusta*). Using the ≥ 50 cut-off, there were one very high-risk horizon species under both BRA and BRA+CCA (green crab *Carcinus maenas*) and 11 additional species under BRA+CCA (extant crown-of-thorns starfish *Acanthaster planci*, white barnacle *Amphibalanus subalbidus*, Pharaoh’s mussel *Brachidontes pharaonis*, upside-down jellyfish *Cassiopea andromeda*, Hepu mitten crab *Eriocheir hepuensis*, Chinese mitten crab *Eriocheir sinensis*, *Hydroides elegans*, giant tiger prawn *Penaeus monodon*, brown mussel *Perna perna*; horizon Australian tubeworm *Ficopomatus enigmaticus* and Manila clam *Ruditapes philippinarum*). Climate-change adjustment increased scores for 27 species (36.5%), resulted in no change for 14 (18.9%), and decreased scores for 33 (44.6%).

*Plants* – ROC analysis yielded an AUC of 0.7738 (95% CI: 0.5574–0.9902) and a calibrated threshold of 31.75 (Fig. 3; Table S5). Based on BRA scores, 13 of 19 species (68.4%) were classified as high risk, and six (31.6%) as medium risk. Under BRA+CCA, 12 species (63.2%) were classified as high risk, and seven (36.8%) as medium risk. Using the ≥ 50 cut-off, there were one very high-risk extant species under both BRA and BRA+CCA (Japanese sea lettuce *Ulva ohnoi*) and two additional species under BRA+CCA (extant shoreline purslane *Sesuvium portulacastrum*, horizon white mangrove *Laguncularia racemosa*). Climate-change adjustment increased scores for eight species (42.1%), resulted in no change for five (26.3%), and decreased scores for six (31.6%).

*Chromists* – ROC analysis yielded an AUC of 0.7047 (95% CI: 0.5548–0.8545) and a calibrated threshold of 30.25 (Fig. 3; Table S5). Based on BRA scores, 36 of 46 species (78.3%) were classified as high risk, and 10 (21.7%) as medium risk. Under BRA+CCA, 36 species (78.3%) were classified as high risk, and 10 (21.7%) as medium risk. Using the ≥ 50 cut-off, there were three very high-risk extant species under both BRA and BRA+CCA (*Lingulaulax polyedra*, golden alga *Prymnesium parvum*, and *Trichodesmium erythraeum*) and 17 additional extant species under BRA+CCA (*Alexandrium minutum*, *Chaetoceros peruvianum*, *Chattonella marina*, *Dinophysis caudata*, *Halamphora coffeiformis*, *Heterosigma akashiwo*, Florida red tide dinoflagellate *Karenia brevis*, *Karenia papilionacea*, *Karenia selliformis*, *Kryptoperidinium triquetrum*, *Margalefidinium polykrikoides*, sea sparkle *Noctiluca scintillans*, *Pseudo-nitzschia delicatissima*, *Pseudo-nitzschia pungens*, *Rhizosolenia delicatula*, *Scrippsiella acuminata*, and

*Tripos furca*). Climate-change adjustment increased scores for 35 species (76.1%), resulted in no change for two (4.3%), and decreased scores for nine (19.6%).

#### 3.3.2 Comparison

Comparison of outcome scores and calibrated risk ranks for the 130 species originally screened and updated in the present study revealed substantive differences (Table S6). These differences arose from both modifications in BRA and BRA+CCA outcome scores and the application of recalibrated risk thresholds. Under the BRA: the risk rank of 104 species (80.0%) remained unchanged. For nine species (7.0%), the rank increased from medium to high; for 12 species (9.2%), it decreased from high to medium; for five species (3.8%), it decreased from medium to low. Under the BRA+CCA: the risk rank of 103 species (79.2%) remained unchanged; for two species (1.5%), it increased from low to medium; for 21 species (16.2%), it increased from medium to high; for three species (2.3%), it decreased from high to medium; for one species (0.8%), it decreased from medium to low.

Comparison with the previous calibration showed numerical differences in thresholds across organismal groups (birds were not previously screened). For fishes, the threshold in the present study was 25.5, compared with previously reported values of 30.50 for brackish fishes and 19.75 for marine fishes. For invertebrates, the threshold was 23.25, whereas earlier values were 26.25 for both brackish and marine categories. For plants, the threshold was 31.75 compared with a previously reported value of 27.50. A threshold of 30.25 was obtained for chromists, a group not evaluated separately in the earlier study. For tunicates, a ROC-derived threshold could not be computed in the present analysis; the previously reported value was 34.25 although with very wide confidence intervals.

#### 3.3.3 Very high-risk species

In total, 39 extant species were ranked as posing a very high risk of invasiveness in the RSA, based on BRA and/or BRA+CCA scores ≥ 50 (Table 2). These species, comprising one bird (2.6% of the total), four fishes (10.3%), three tunicates (7.7%), nine invertebrates (23.1%), two plants (5.1%), and 20 chromists (51.3%), were therefore identified for progression to comprehensive risk assessment and subsequent management action within the regional biosecurity framework. At the Member State level, the distribution of very high-risk extant species identified for prioritization was uneven: five species were relevant to Bahrain, 20 to Iran, 10 to Iraq, 26 to Kuwait, 21 to Oman, six to Qatar, eight to Saudi Arabia, and 12 to the United Arab Emirates.

**Table 2.**
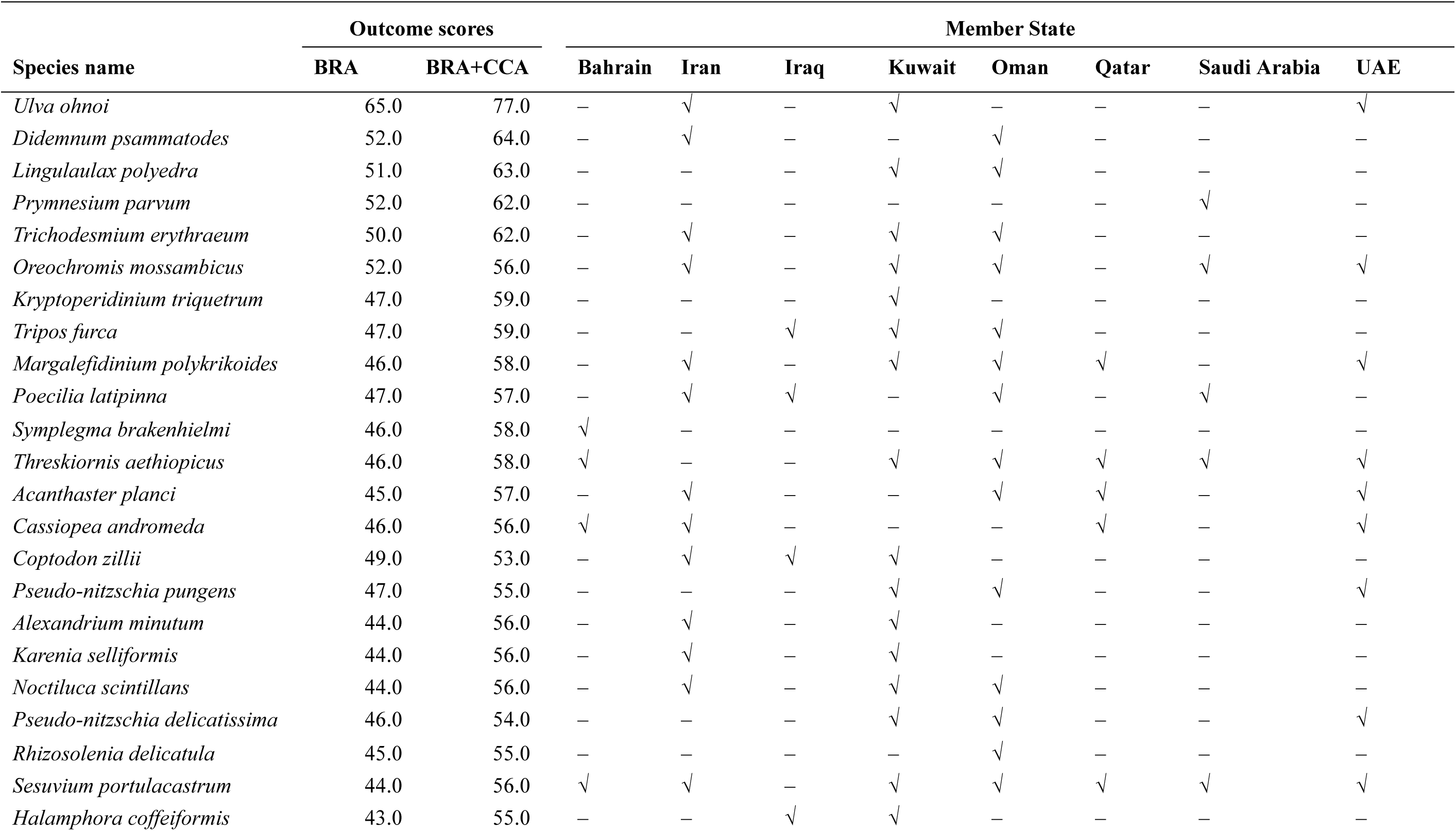
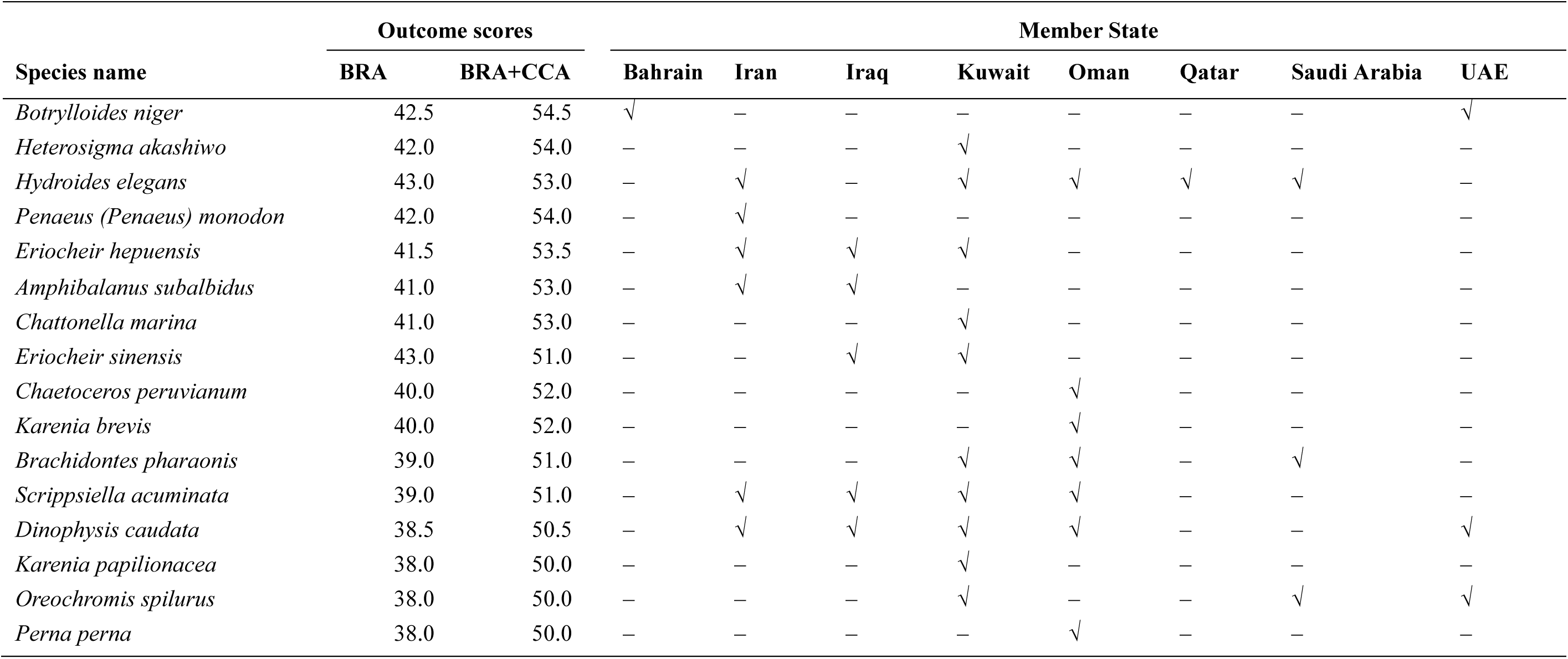
Extant species assessed with the Aquatic Species Invasiveness Screening Kit for the RSA and ranked as carrying a very high risk of invasiveness with indication of their reporting in each of the eight ROPME Member States. BRA = Basic Risk Assessment outcome score; BRA+CCA = BRA + Climate Change Assessment outcome score. Species are listed in descending order from higher to lower sum of BRA and BRA+CCA. See also Fig. 2.

In total, eight horizon species were ranked as carrying a very high risk of invasiveness in the RSA, based on BRA and/or BRA+CCA scores ≥ 50 (Table 3). These species, These species, comprising three fishes (37.5%), one tunicate (12.5%), three invertebrates (37.5%), and one plant (12.5%), were identified for progression to comprehensive risk assessment and subsequent risk management within the regional biosecurity process, consistent with their classification under the very high-risk category.

**Table 3.**
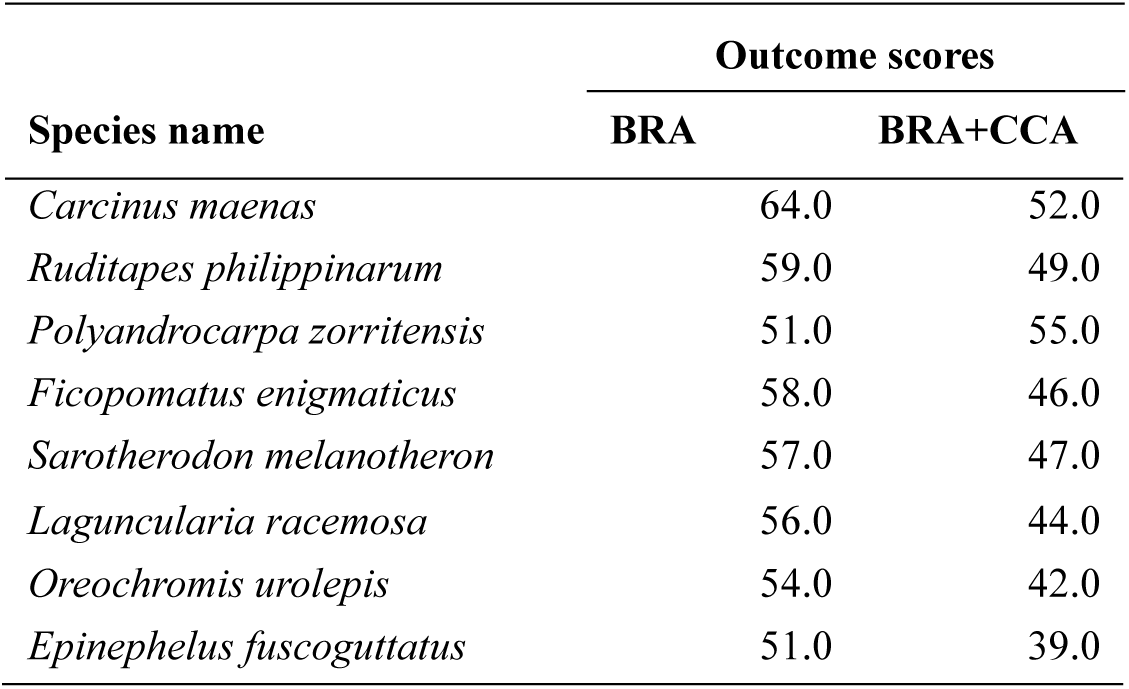
Horizon species screened with the AS-ISK for the RSA and ranked as carrying a very high risk of invasiveness. Outcome scores are reported as per Table 2.

## 4 Discussion

This study establishes the first harmonized and calibrated regional biosecurity baseline for the RSA by integrating Member State-level validated categorization of extant species status within a novel, data-intensive framework with systematic risk identification of both extant and horizon species under present and projected climate conditions, implemented through a globally calibrated decision-support tool (Vilizzi et al., 2021, 2025a). Beyond revising and consolidating species inventories within a robust and evidence-based analytical structure, the results reveal coherent ecological patterns shaped by the extreme environmental conditions and intensifying human-mediated pressures characteristic of the RSA, consistent with documented invasion dynamics in semi-enclosed and heavily impacted marine systems (Occhipinti-Ambrogi and Savini, 2003; Rilov and Crooks, 2009; Ojaveer et al., 2017, 2018; Tiralongo et al., 2022). Collectively, this framework provides an operational foundation for anticipatory and transboundary marine biosecurity in the RSA, aligned with the adoption of best-practice principles in invasion science (Hewitt and Campbell, 2007; Cook et al., 2016; Campbell and Hewitt, 2025; Jayachandran et al., 2026), and establishes a structured pathway toward full risk assessment and subsequent risk management of species identified as high priority (i.e. very high risk) (Vilizzi et al., 2022a).

### 4.1 Ecological patterns and invasion dynamics

The taxonomic structure of the extant species dataset reflects the environmental singularity of the RSA (Sheppard et al., 2010; Naser, 2014; Lincoln et al., 2021). In such extreme marine ecosystems, harsh physicochemical conditions constrain establishment by many species while favouring stress-tolerant organisms pre-adapted to high temperature variability and fluctuating salinity (Price et al., 1993; Vaughan et al., 2019; Torquato, 2025). Invasion patterns in the RSA are therefore best interpreted mechanistically through trait–environment matching operating alongside strong propagule pressure and human-induced habitat modification (Kolar and Lodge, 2001; Ojaveer et al., 2018).

Climate-adjusted risk scores indicate that projected warming may further modify this selective landscape (Rahel and Olden, 2008; Hellmann et al., 2008). Increases in risk under the climate-change scenario were particularly evident among chromists, consistent with the thermal sensitivity and opportunistic dynamics of many bloom-forming species (Hallegraeff, 2010; Wells et al., 2015; D’Souza, 2022). In contrast, the other organismal groups, exhibited offsetting increases and decreases in risk, producing no uniform directional response. Given that baseline thermal regimes in the RSA already approach upper physiological tolerance limits for many species, further warming is likely to redistribute invasion performance rather than uniformly amplify risk (Walther et al., 2009; Sorte et al., 2010; Clarke et al., 2020).

Within this environmental and climatic context, the designation of 39 extant and eight horizon species as very high risk demonstrates the functional breadth of invasion exposure within the RSA and indicates that invasion pressure is systemic rather than taxonomically constrained. Their designation as very high risk reflects convergence across environmental matching, pathway alignment, invasion history, and documented or strongly inferred ecological and socio-economic impacts, consistent with established invasion theory linking species traits, propagule pressure, and recipient-environment characteristics (Catford et al., 2009; Lockwood et al., 2005; Blackburn et al., 2011).

A substantial component of the very high-risk extant species comprises bloom-forming chromists, including multiple toxin-producing dinoflagellates and diatoms associated with HABs (Table 2). Their prominence in the Inner subregion is consistent with the RSA’s semi-enclosed hydrography, intense maritime connectivity, expanding aquaculture, and nutrient-enriched coastal zones (e.g. Al-Yamani et al., 2012; Al Shehhi et al., 2014; Al-Azri et al., 2015). Additional opportunistic bloom-formers and primary producers include *Noctiluca scintillans*, *Sesuvium portulacastrum*, *Trichodesmium erythraeum*, and *Ulva ohnoi*, all capable of rapid biomass accumulation under eutrophic conditions and indicative of nutrient-mediated regime shifts (Hirahoka et al., 2004; Al-Yamani et al., 2012; Al-Azri et al., 2015; Patzelt and Lupton, 2021). Collectively, these species present acute risks to fisheries, aquaculture, desalination infrastructure, and public health through toxin production, fish mortality events, and hypoxia following bloom decay (Lincoln et al., 2021).

Other very high-risk extant species include benthic ecosystem engineers and fouling organisms such as the tunicates *Botrylloides niger*, *Didemnum psammatodes*, and *Symplegma brakenhielmi* (Sheets et al., 2016; Lambert, 2019; Mohaddasi et al. 2019), the serpulid *Hydroides elegans* (Nedved and Hadfield, 2008), the barnacle *Amphibalanus subalbidus* (Yasser et al., 2022), and the bivalves *Brachidontes pharaonis* and *Perna perna* (Borthagaray and Carranza, 2007; Battiata et al., 2024) – all closely associated with shipping and port environments and capable of increasing infrastructure costs while restructuring native benthic assemblages. Mobile crustaceans and commercially relevant decapods include *Eriocheir hepuensis*, *Eriocheir sinensis*, and *Penaeus (Penaeus) monodon*, which have demonstrated capacity to modify sediment structure, alter trophic interactions, and affect aquaculture systems (Naser et al., 2011; Ferreira et al., 2025). Higher consumers include the invertebrates *Acanthaster planci* and *Cassiopea andromeda* (Nabipour et al., 2015; Uthicke et al., 2024), the fishes *Coptodon zillii*, *Oreochromis mossambicus*, *Oreochromis spilurus*, and *Poecilia latipinna* (Russell et al., 2012; Mohamed and Al-Wan, 2020; U.S. Fish and Wildlife Service, 2020; Güçlü et al., 2021), and the bird *Threskiornis aethiopicus* – species capable of restructuring food webs and exerting strong competitive or predatory pressures (Volponi et al., 2025).

The eight horizon species reinforce this pattern. They encompass the globally invasive benthic engineers and fouling invertebrate *Ficopomatus enigmaticus* and the tunicate *Polyandrocarpa zorritensis* (Charles et al., 2018; Stabili et al., 2015), the highly dispersive decapod *Carcinus maenas* (Ens et al., 2022), the commercially valuable bivalve *Ruditapes philippinarum* (Coelho et al., 2021), the fishes linked to aquaculture and trade pathways *Epinephelus fuscoguttatus Oreochromis urolepis*, and *Sarotherodon melanotheron* (Cassemiro et al., 2018; Chaianunporn et al., 2024; Zhang et al., 2024), as well as habitat-forming coastal vegetation (*Laguncularia racemosa*: Zhang et al., 2025). Their ranking as very high risk despite current absence reflects strong environmental matching and clear alignment with dominant introduction pathways, indicating credible establishment potential under plausible propagule-pressure scenarios.

### 4.2 Status validation and threshold recalibration

Re-evaluation of the species originally screened showed that approximately 18% required status reassignment, highlighting how even formally curated databases can yield divergent classifications when re-evaluated under validated criteria. In addition, re-examination of the published literature revealed that the distribution of species across Member States previously reported was frequently inaccurate. A further distinction concerns the evidentiary basis and spatial scope of the screenings. Whereas the original assessments were conducted primarily by researchers external to the ROPME region and were limited to the Inner and Middle RSA, the present re-evaluation incorporated systematic input and validation from in-country experts across Member States and encompassed the entire RSA (i.e. Inner, Middle, and Outer subregions).

The above discrepancies underscore the need for rigorous, region-specific validation of databases of ‘non-native’ species, particularly in marine systems where historical baselines, cryptogenic status, and shifting biogeographic boundaries often blur classification criteria (e.g. Glamuzina et al., 2024). Misclassification of extant, horizon, and native species, together with inaccurate Member State distribution records, has direct consequences for the establishment of biosecurity baselines, especially in highly impacted ecosystems such as the RSA. Accordingly, systematic scrutiny of pre-compiled non-native species lists, including verification of national-level occurrence data, should constitute a mandatory quality-control step within risk analysis frameworks (Vilizzi and Piria, 2022). Database-driven screenings, therefore, provide essential starting points, particularly in data-heterogeneous systems, but they remain provisional unless systematically reconciled with primary literature and validated national-level evidence (Pagad et al., 2018; Briski et al., 2024; Gómez-Suárez et al., 2025).

A key refinement introduced in the present study concerns the categorization of extant species status. Our systematic re-evaluation has demonstrated that a substantial proportion of the original species (*n* = 28; 23%) are native to the RSA and to at least one Member State, but exhibit nuisance characteristics under present environmental conditions, as reflected in their categorization as nuisance native. The same argument holds for cryptogenic species, whose biogeographic origin cannot be resolved with confidence due to historical data gaps, early introductions, or natural range expansions predating systematic monitoring (Carlton, 1996; Jarić et al., 2019). The uneven proportional distribution of native-related, nuisance, and non-native categories across Member States illustrates structural asymmetries in invasion pressure, ecological impact expression, and documentation intensity. This hierarchical and impact-refined species status categorization ensures transparent differentiation between origin and impact status, facilitates progression through the full risk analysis process (risk identification, risk assessment, and risk management), and supports harmonized regional biosecurity reporting across ROPME Member States.

Comparison with Clarke et al. (2020) highlights the influence of both habitat structuring and dataset composition on ROC-derived threshold calibration. Clarke et al. (2020) computed separate BRA thresholds for brackish and marine fishes and invertebrates within the Inner and Middle RSA, whereas the present study intentionally pooled brackish and marine species for fishes, invertebrates, and plants to reflect ecological connectivity and region-wide management integration across the RSA. In addition, the present analysis screened 60 additional species, thereby expanding and restructuring the underlying score distribution. The intermediate fish threshold and reduced invertebrate threshold observed here therefore represent recalibration under pooled habitat structure and expanded species coverage rather than methodological divergence. The higher plant threshold similarly reflects optimisation within a broader and more taxonomically inclusive dataset. These contrasts underscore that screening thresholds are dataset-dependent analytical parameters rather than fixed biological constants.

The recalibration of thresholds in the present study further reinforces the dynamic nature of non-native species risk analysis, which should adopt a review-and-revision approach concerning both the risk analysis process and the management strategy for the species of concern (Mumford et al., 2010). As demonstrated by re-screening exercises (Tarkan et al., 2017; Yoğurtçuoğlu et al., 2025), periodic updating of risk screenings can result in substantial changes in species categorization and risk ranking due to newly available information, methodological refinement, and incorporation of climate change projections. This underscores that risk analysis is not a static classification exercise but an iterative process requiring structured review and recalibration. In the context of the RSA, where environmental conditions are rapidly changing and maritime connectivity remains intense, maintaining updated and regionally calibrated thresholds is essential to ensure that prioritization reflects current ecological realities rather than historical data snapshots. Regular revision of screenings therefore constitutes a core component of adaptive biosecurity governance under the ROPME framework.

By integrating multi-source cross-verification, taxonomic harmonization, and explicit status categorization, the present study transforms exploratory screening into a reproducible and defensible analytical platform (Leung et al., 2012; Roy et al., 2019). Misclassification of species origin or invasion status is not trivial. Overestimation of non-native occurrence, for example through the inclusion of native or nuisance species within non-native inventories, inflates perceived regional impacts and distorts cross-jurisdictional comparisons. Conversely, underestimation, such as when extant species are erroneously treated as horizon taxa, delays recognition of realised invasion pressure and postpones appropriate management responses (Latombe et al., 2017; Ricciardi et al., 2017). Once incorporated into global databases, such errors can propagate across subsequent studies and policy instruments (McGeoch et al., 2012; Pagad et al., 2022). The systematic differentiation among status categories implemented in our study improves ecological interpretability and reduces conflation of distinct invasion processes (Carlton, 1996; Blackburn et al., 2011).

### 4.3 Establishing a biosecurity baseline

Explicit differentiation between extant and horizon species strengthens the anticipatory dimension of regional invasion science (Vilizzi et al., 2022a). Horizon scanning, an important requirement in non-native species risk identification studies (e.g. (Clarke et al., 2020; O’Shaughnessy et al., 2023; Glamuzina et al., 2024), when grounded in documented invasion histories, pathway analysis, and environmental suitability rather than speculative listing, enables structured anticipation of plausible future establishments. Similarly, the inclusion of non-native captive species as extant further acknowledges aquaculture, ornamental trade, and confinement as latent introduction pathways.

Marine biological invasions in semi-enclosed seas are inherently transboundary, shaped by shared circulation systems, intense maritime connectivity, and highly mobile introduction vectors (e.g. Katsanevakis et al., 2014; Giangrande et al., 2020; Tiralongo et al., 2022). Uneven documentation and monitoring capacity among Member States can generate informational asymmetries that hinder coordinated response and effective management. A harmonized regional baseline reduces these asymmetries by providing a standardized evidential reference for surveillance prioritization, reporting comparability, and risk progression within cooperative governance frameworks (e.g. Tsiamis et al., 2019; Figuereido et al., 2024).

As emphasized, this study addresses the first stage of the risk analysis process, namely risk identification (Vilizzi et al., 2022a). Progression to full risk assessment of the very high-risk species will enable quantitative evaluation of impact magnitude, ecological interactions, and socio-economic implications, followed by structured risk management within established biosecurity frameworks (Leung et al., 2012). Although species classified as high risk also warrant continued monitoring and periodic re-evaluation, progression to full risk assessment is prioritized for very high-risk species in recognition of the substantial technical, financial, and institutional resources required for comprehensive assessment and subsequent management implementation (Copp et al., 2016a, 2016b). Full risk assessment entails detailed pathway analysis, ecological modelling, stakeholder consultation, and potential regulatory intervention, which impose considerable demands on national authorities. In a transboundary governance context such as the RSA, where capacity and resources vary among Member States, prioritizing very high-risk species represents a pragmatic and proportionate allocation strategy that maximizes regional management effectiveness while maintaining precautionary oversight of other high-risk species. Establishing a calibrated and transparent baseline is therefore a prerequisite for consistent prioritization and policy-relevant decision-making (Roy et al., 2019, 2024).

The analytical framework applied here, combining taxonomic standardization, Member State-level reconciliation, horizon integration, and ROC-based calibration, is transferable to other marine systems characterized by high connectivity and transboundary governance (e.g. Katsanevakis et al., 2014; Tsiamis et al., 2019). Structured and transparent risk-screening approaches enhance consistency, comparability, and reproducibility across jurisdictions and taxonomic groups (e.g. Wei et al., 2021; Mumladze et al., 2022; Glamuzina et al., 2024). thereby facilitating operational integration into national environmental management plans and regional policy instruments.

### 4.4 Structural uncertainty and regional monitoring heterogeneity

As with all regional baselines, the present assessment reflects underlying heterogeneity in taxonomic resolution, molecular verification, monitoring intensity, and national documentation across Member States. These dimensions constitute structured sources of uncertainty inherent to transboundary invasion assessments and require continued refinement through coordinated regional action. Addressing such variability reinforces the need for harmonized monitoring protocols, dynamically curated species inventories, and integrated marine data systems under the ROPME Strategic Directions 2026–2030 (AlDimashki & AlAhmad, 2025).

The occurrence of species complexes and unresolved synonymies complicates status assignment. Examples in this study include the recognition in the Gulf of *Acanthopagrus arabicus* sp. nov. and *A. sheim* as distinct from *A. latus* (Iwatsuki, 2013), the differentiation of *Acanthaster planci* and *A. mauritientis* in Oman and the UAE (Uthicke et al., 2024), and the continuing taxonomic uncertainty surrounding *Brachidontes pharaonis* (Battiata et al., 2024).

Records of Indo-Pacific species such as *Actinotrichia fragilis*, *Dichotomaria marginata*, *Hypnea musciformis*, *Izziella orientalis*, and *Trichogloea requienii* (Ibrahim, 2018), classified here as neonative in some Member States, would benefit from molecular corroboration to exclude the possibility that their apparent recent occurrence reflects historical under-detection rather than genuine range shifts within the RSA. The incorporation of molecular tools, including DNA barcoding and environmental DNA approaches, into routine monitoring frameworks would strengthen confidence in status assignments and enhance early detection of emerging introductions (e.g. Tarkan et al., 2022).

Monitoring and reporting efforts in the RSA, as elsewhere, tend to favour conspicuous, economically important, or ecologically impactful organisms such as fishes, decapods, and harmful algal bloom-forming microalgae, whereas smaller-bodied, cryptic, or taxonomically challenging groups, including meiofauna, benthic invertebrates, and certain planktonic assemblages, remain comparatively underrepresented (Lincoln et al., 2021). Such imbalance may skew apparent invasion patterns and underestimate diversity within understudied taxa. Targeted investment in underrepresented taxonomic groups, including standardized benthic surveys, plankton monitoring, and capacity building in specialist taxonomy, would help reduce detection bias and refine regional risk prioritization.

Finally, of the eight Member States, national lists of invasive species are currently available for Iran (Department of Environment, Islamic Republic of Iran, 2025), Iraq (Ministry of Health and Environment, Republic of Iraq, 2018), Kuwait (Al-Yamani, 2021), and the UAE (UAE MOCCAE, 2022), with an additional list for Saudi Arabia in preparation (E. I. Sabbagh, unpublished data); no national lists are yet available for Bahrain, Oman, or Qatar. This heterogeneity in baseline documentation inevitably influences detectability and reporting consistency across the region. Establishing standardized national inventories for all Member States, coupled with periodic regional synthesis updates, would substantially improve comparability and early-warning capacity.

## 5 Conclusion

Beyond methodological refinement, the present recalibration represents a structural advancement in regional biosecurity governance under the ROPME framework. By aligning status categorization, species inventories, and ROC-derived thresholds with the full spatial unit of governance and embedding systematic in-country expert validation into the screening process, the framework reduces jurisdictional inconsistencies and strengthens decision legitimacy across Member States. In an environmentally extreme and highly connected marine system such as the RSA, validated and governance-aligned risk identification provides the evidential foundation for progression to full risk assessment and coordinated biosecurity management of species ranked as (very) high risk. The framework is explicitly designed to accommodate new data and periodic recalibration, ensuring adaptive updating while preserving analytical coherence and comparability across assessment cycles. By integrating ecological insight, expert validation, and governance-scale calibration within a reproducible analytical architecture, the RSA serves not only as a case study of biological invasion under extreme conditions, but also as a transferable model for harmonized regional biosecurity governance in other semi-enclosed and climatically stressed marine systems.

## CRediT authorship contribution statement

**Lorenzo Vilizzi**: Conceptualization, Methodology, Formal analysis, Writing – original draft, Validation, Writing – review & editing, Supervision. **Ali Mansoor Abbas**: Validation, Writing – review & editing. **Maryam A. Mubarak**: Validation, Writing – review & editing. **Mohammad Hadi Alavi**: Validation, Writing – review & editing. **Mahtab Shojaei**: Validation, Writing – review & editing. **Daryoush Moghaddas**: Validation, Writing – review & editing. **Hossein Rahmani**: Validation, Writing – review & editing. **Ahmed A. R. Albu Salih**: Validation, Writing – review & editing. **Mohammed F. A. Al-Khayyat**: Validation, Writing – review & editing. **Abbas J. Al-Faisal**: Validation, Writing – review & editing. **Aisha F. Al-Marhoun**: Validation, Writing – review & editing. **Ali H. Abdulhussain**: Validation, Writing – review & editing. **Jumana Alkhamees**: Validation, Writing – review & editing. **Qusaie E. Karam**: Validation, Writing – review & editing. **Wafa’a Behbehani**: Validation, Writing – review & editing. **Mohammed Al Rezaiqi**: Validation, Writing – review & editing. **Muna Al Tarshi**: Validation, Writing – review & editing. **Salman F. Al-Dosari**: Validation, Writing – review & editing. **Amal M. Al Jamaei**: Validation, Writing – review & editing. **Mohammed Elnaiem A. El Mahdi**: Validation, Writing – review & editing. **Abdirahman A. Mohamed**: Validation, Writing – review & editing. **Eman I. Sabbagh**: Validation, Writing – review & editing. **Ebrahim A. Jamali**: Validation, Writing – review & editing. **Nahla Mezhoud**: Validation, Writing – review & editing. **Obaid A. H. Al Shamsi**: Validation, Writing – review & editing. **Zainab Al-Wazzan**: Conceptualization, Validation, Writing – review & editing, Supervision.

## Acknowledgments

This study was conducted within the framework of a regionally coordinated programme on marine invasive species led by the Regional Organization for the Protection of the Marine Environment (ROPME). ROPME is gratefully acknowledged for its scientific coordination, logistical support, and facilitation of collaboration among the Member States of the ROPME Sea Area. The views expressed in this article are those of the authors and do not necessarily reflect the official positions of ROPME or its Member States.

